# A hierarchical generative model reveals enhanced latent precision of brain-body interaction dynamics during interoceptive attention

**DOI:** 10.64898/2026.04.05.716599

**Authors:** Kazushi Shinagawa, Hayato Idei, Satoshi Umeda, Yuichi Yamashita

## Abstract

Brain–body interactions (BBIs) are fundamental to cognition and mental health, but their continuous multimodal dynamics remain difficult to extract. Previous approaches have been largely observational, and few frameworks enable these interacting processes to be modeled within an integrated generative system. Here, we applied a Predictive-Coding-Inspired Variational RNN (PV-RNN) to simultaneous EEG, ECG, and respiration recordings obtained from 33 participants during exteroceptive and interoceptive attention. The model learned a temporal hierarchy spanning modality-specific dynamics, multimodal associative integration, and sequence-level global states, and accurately reconstructed unseen physiological sequences. Specifically, the intermediate associative layer successfully captured the core complexities of BBI by extracting multiscale, nonlinear, and bidirectional coupling dynamics with variable temporal lags. Furthermore, the estimated precision (inverse variance) of latent variables representing BBI dynamics within this multimodal associative layer increased significantly during interoceptive attention. The magnitude of this condition-dependent precision enhancement correlated positively with subjective adaptive body controllability and negatively with psychiatric vulnerabilities, including rumination and trait anxiety. These findings identify a latent physiological signature of interoceptive attention and establish hierarchical generative modeling as an interpretable framework for extracting continuous BBI dynamics and linking multimodal physiology to cognitive and clinical characteristics.

## Introduction

Brain–body interaction (BBI) has become a central topic in cognitive neuroscience because cognition is increasingly understood as a process shaped by ongoing interactions between neural and bodily activity (Buzsáki & Tingley, 2023; Ribeiro & Oliveira-Maia, 2025). Physiological dynamics in visceral signals and their neural processing influence a diverse range of cognitive functions, including attention (Petzschner et al., 2019), emotion (Critchley & Garfinkel, 2017), memory (Messina et al., 2022; Umeda et al., 2016), decision-making (Fujimoto et al., 2021; Herman et al., 2021), self-representation (Babo-Rebelo et al., 2019), thought (Ito et al., 2019; Sakuragi et al., 2024), and behavior (Johannknecht & Kayser, 2022; Palser et al., 2021), and their dysregulation has been implicated in mental health and psychiatric vulnerability (Gregorio & Battaglia, 2024; Khalsa et al., 2018). Despite this importance, current approaches often capture only partial aspects of BBI, reducing the brain–body system to isolated indices rather than treating it as an integrated dynamical process.

Existing approaches have provided important but still fragmentary insights. Event-based methods, such as heartbeat-evoked or respiration-locked neural responses, have greatly advanced interoception research by isolating afferent processes tied to specific physiological events (Poppa et al., 2022; Schandry et al., 1986; von Leupoldt et al., 2011). For instance, these methods have demonstrated that cortical responses to individual heartbeats are modulated by interoceptive attention (Petzschner et al., 2019). However, these approaches reduce continuous multimodal dynamics to discrete events and are less suited for describing ongoing interactions across bodily systems. Time-series approaches, including mutual information and transfer entropy (Antonacci et al., 2020; Martínez Vásquez et al., 2023; Pernice et al., 2019), have overcome the need for event segmentation, revealing how directed information flow between the brain and body is modulated by changes in cognitive and affective states. While these metrics quantify directed information flow, their reliance on static pairwise summaries limits their ability to capture continuous and state-dependent nonlinear dynamics. Although a recent model incorporates physiological constraints to address non-linearity and bidirectionality, it remains limited by pre-specified frequency bands and difficulties in extending to diverse modalities (Candia-Rivera, 2023).

These limitations arise in part from the structure of BBI itself. Neural activity exhibits fast and irregular fluctuations, whereas cardiac and respiratory signals show slower rhythms, event-like morphologies, and variable timing. Their interactions are nonlinear, distributed across multiple timescales, and often involve bidirectional influences with unknown or variable lags (Candia-Rivera, 2022; Lin et al., 2016; Schiecke et al., 2019). Given this complexity, conventional measures of brain–body coupling that focus solely on statistical dependencies among observed signals leave it unresolved whether the observed relationships arise from direct interactions, temporally lagged influences, or common latent drivers. Although recent multimodal deep-learning methods offer greater flexibility for integrating heterogeneous physiological data, they have largely been developed for multimodal fusion and downstream predictive tasks rather than for modeling the latent dynamical structure of those interactions (Kline et al., 2022; Pillalamarri & Shanmugam, 2025). A generative framework could address these limitations by modeling the hidden dynamical structure through which these signals are co-generated over time, allowing BBI to be examined as an inferred latent interaction process rather than only through its observable traces (Durstewitz et al., 2023). From these perspectives, a useful generative model of BBI should 1) preserve modality-specific multiscale structure, 2) capture bidirectional and nonlinear cross-modal coupling with temporal lags, 3) integrate time-varying uncertainty for state-dependent and modality-specific fluctuations, and 4) yield interpretable latent representations for application to cognition and mental health.

Here, we address this gap using a predictive-coding-inspired variational recurrent neural network (PV-RNN) trained on simultaneous EEG, ECG, and respiration recorded during focused attention on external or internal stimuli (Shinagawa et al., 2025). PV-RNN combines deterministic and stochastic latent variables within a temporal hierarchy (Ahmadi & Tani, 2019; Idei et al., 2022; Takahashi et al., 2025), providing the representational capacity to model BBI, where interactions unfold across multiple timescales, include substantial noise and variability, and depend on probabilistic coupling across modalities. Recent work has shown that this model family can generate highly stochastic physiological signals while extracting underlying channel-to-channel associations and global states, such as awake and anesthetized conditions, in a data-driven manner (Takahashi et al., 2025). Furthermore, the model has been shown to extract cross-modal associative structures and global mode transitions through hierarchical latent representations and dynamically modulated precision weighting (Idei et al., 2022). Building on these findings, we apply a hierarchical PV-RNN to human physiological signals to examine whether BBI can be described within an integrated generative framework, where modality-specific lower levels extract within-signal dynamics, an intermediate associative level integrates slower cross-modal interactions, and a higher-level state captures global sequence-level context.

We evaluate the model in a stepwise manner that aligns with the generative requirements of BBI discussed above. First, we assess whether PV-RNN can accurately reconstruct multimodal data at the waveform level preserve physiologically meaningful structures. Second, building on this generative capability, we examine whether the intermediate associative layer captures multimodal coupling. Specifically, using ablation and dynamical analyses, we examine how these shared latent variables contribute to multimodal generation and assess whether their dynamics are consistent with key aspects of BBI, including multiscale dynamics, nonlinear interactions, bidirectionality, and temporal lags. Third, we evaluate the construct validity of the extracted latent states by assessing their correspondence with experimentally manipulated attentional conditions and known physiological coupling patterns. Finally, we investigate whether the highest-level latent representations spontaneously segregate global sequence-level information, such as distinct attentional states and individual identity. By framing BBI as hierarchical multimodal generation, this study offers a computational framework for moving beyond fragmented physiological indices.

## Results

### Datasets and Model Overview

#### Training and Test

To establish a computational framework for investigating BBI, we required a dataset providing simultaneous recordings of brain activity and autonomic signals. Furthermore, to enable evaluation of the extracted BBI dynamics, the dataset needed to explicitly encompass distinct cognitive states in which dynamic modulations of BBI are theoretically expected. Therefore, we focused on how attentional allocation is modulated between the external environment and internal bodily sensations. We trained the PV-RNN using a dataset comprising the simultaneous EEG, ECG, and respiratory (RESP) recordings acquired during exteroceptive (Sound Focus; SF) and interoceptive (Breath Focus; BF) attention tasks (Shinagawa et al., 2025). In SF, participants responded as quickly as possible to randomly presented pure tones (inter-tone interval: 1.2–2.0 sec), whereas in BF they responded at the end of each exhalation; the same pure tones were also presented in BF to equate the auditory environment. In both conditions, participants performed self-reports when they noticed their attention had drifted away. We extracted the periods 1–5 s preceding these reports, representing intervals when participants were presumably focused on the task (i.e., on sounds or their breath). Each sequence was 4 sec long sampled at 100 Hz (400 time points) and comprised 19 EEG channels, 2 RESP channels, and 1 ECG channel; the 19 EEG channels were selected to broadly cover the scalp.

The model was trained for multimodal generation on 660 sequences from 33 participants (10 SF and 10 BF per participant). After training, to verify that the capability of multimodal data generation is not driven by overfitting on training datasets, we fixed the synaptic weights and evaluated whether accurate reconstruction could be achieved solely by inferring the latent state. This generalization was tested on two distinct datasets: held-out sequences from trained participants (cohort test for within-subject generalization; n = 30, 180 sequences), and sequences from completely novel participants (unobserved participant test for across-subject generalization; n = 9, 54 sequences). To ensure comparability, both test sets were balanced to contain exactly three SF and three BF sequences per participant, excluding individuals with insufficient usable data.

#### System Overview

The model has a hierarchical spatiotemporal architecture (Fig. 1a) (Ahmadi & Tani, 2019; Idei et al., 2022; Takahashi et al., 2025; Yamashita & Tani, 2008) designed to represent the specific characteristics of the dataset across three levels: modality-specific modules at the lowest level capturing the independent, fast dynamics of each signal (e.g., ECG, Resp, EEG), a multimodal-associative level integrating shared cross-modal information at intermediate timescales, and a global state level capturing sequence-level contexts, such as individual- or task-related overall dynamic changes, at the highest level. In the present study, latent variables in the global state layer were treated as sequence-specific constants (i.e., fixed within each sequence and not updated over time), so that they encode stable, sequence-level context while moment-to-moment dynamics are captured by lower levels. Dynamics at each level are implemented by stochastic latent variables (*z*(*l*)), which encode level-specific uncertainty, together with deterministic states (*d*(*l*)), which carry timescale-dependent recurrent dynamics.

**Fig. 1.**
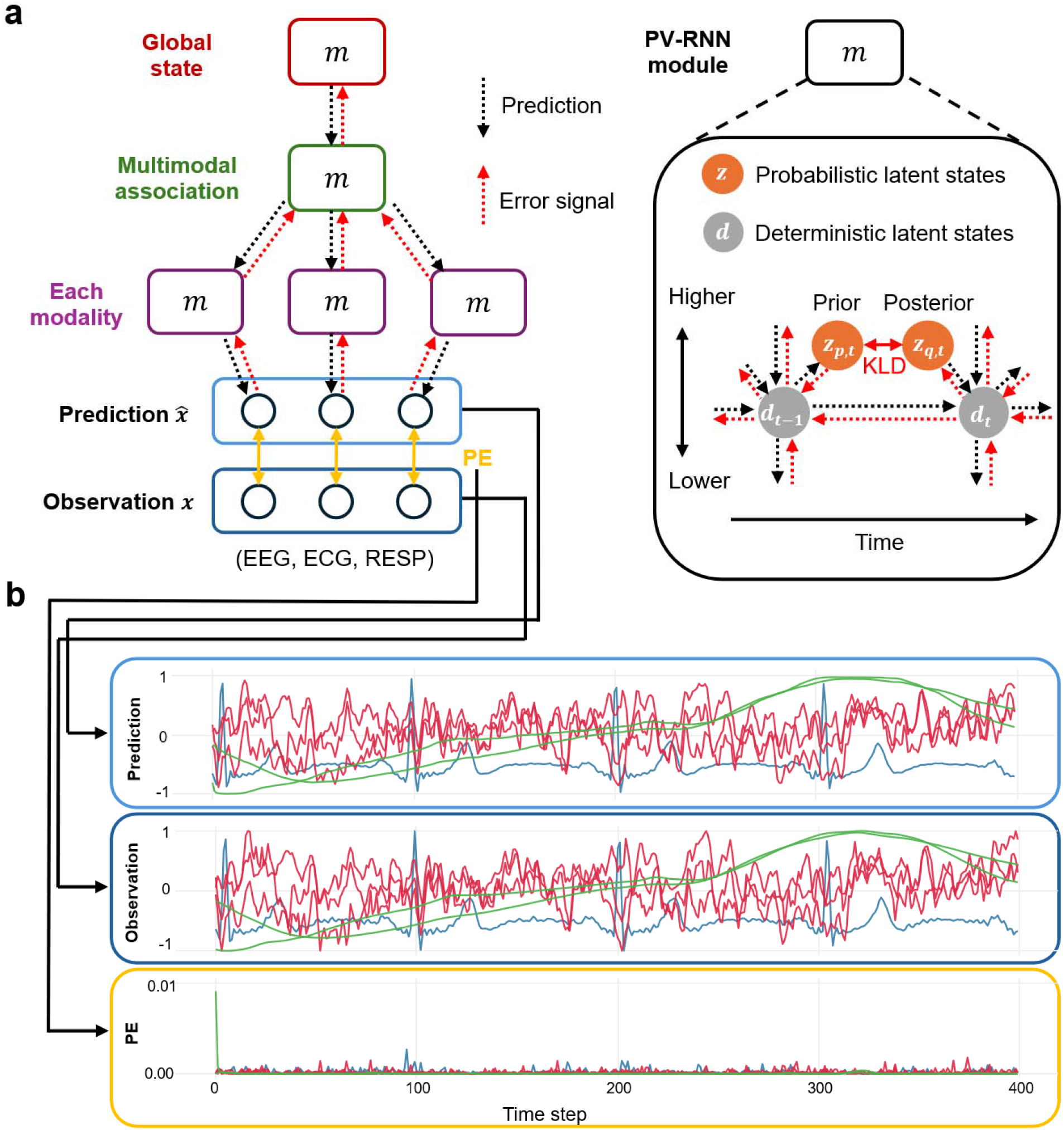
Predictive-coding-inspired variational recurrent neural network. **a**. Structure and information processing of Predictive-coding-inspired variational recurrent neural network (PV-RNN). PE: prediction error. KLD: Kullback-Leibler divergence. **b**. Examples of prediction and observed data during learning. EEG signals are visualized by selecting 3 channels from the original channels for visual clarity. The red, blue, and green lines represent EEG, ECG, and respiratory signals, respectively. Three EEG channels are selected for visual clarity.

PV-RNN is trained as a dynamic generative model (Fig. 1b). During the training phase, model parameters are optimized across sequences, while latent states are inferred to explain the observed multimodal time series. Given an observed sequence, latent states are initialized from the prior and used to generate top-down predictions for each modality. The discrepancy between predicted and observed signals is then used to iteratively refine latent states and synaptic weights, yielding posterior latent states that better explain the observed time series. In the test phase, all model parameters were held fixed, and only latent variables were optimized to reduce prediction error for each sequence, yielding posterior latent variables that better explain the observed time series. Additional details of this inference scheme are provided in Supplementary Fig. 1-2 and Methods.

### PV-RNN accurately reconstructs multimodal EEG–ECG–respiratory time series

First, we assessed whether the trained PV-RNN accurately reconstructs multimodal data at the waveform level and preserves physiologically meaningful structures. Following training, the PV-RNN learned a stable generative model of multimodal dynamics across EEG, ECG, and RESP signals in both BF and SF conditions. In both datasets, the reconstructed signals appeared physiologically plausible on visual inspection. Fig. 2a shows representative reconstructed sequences of unobserved datasets, illustrating that the model captures diverse temporal structures across modalities. The trained model accurately reconstructs both fast EEG fluctuations and slower respiratory dynamics within a single hierarchical generative model. In addition, it reproduces event-like ECG morphology with a stereotyped waveform shape while allowing variability in event timing, consistent with the quasi-deterministic shape and stochastic occurrence of cardiac cycles. Remarkably, prediction errors for unobserved participants were as low as those for within-cohort held-out sequences, with error distributions showing substantial overlap at small magnitudes (Fig. 2b). These results suggest that the model captured multimodal physiological data faithfully at the waveform level and that downstream inferences on latent states and their associations are unlikely to be driven by overfitting or idiosyncratic properties of the training datasets.

**Fig. 2.**
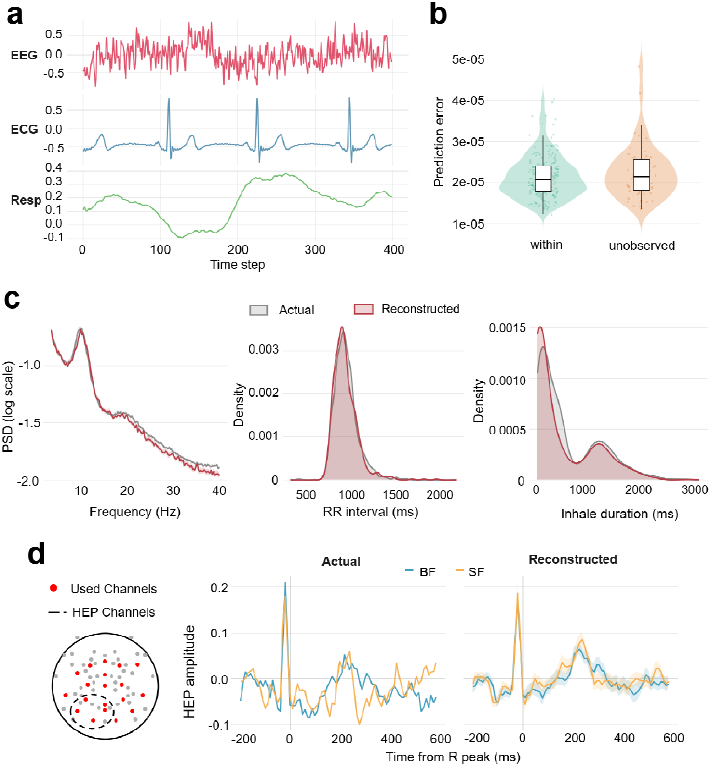
Reconstructed multimodal signals and physiological properties. **a**. Examples of prediction for datasets from unobserved participants. EEG signals are visualized by selecting 1 channel from the original channels for visual clarity. **b**. Distributions of mean prediction error of each sequence across held-out within-cohort and unobserved participants. **c**. Comparison of summary physiological properties between the actual physiological signals from the training datasets and reconstructed data pooled across within-cohort and unobserved test sequences. Left, mean EEG power spectral density across channels. Middle, distribution of mean RR intervals. Right, distribution of inhalation-phase duration within sequences. **d**. Heartbeat-evoked potential in the actual and reconstructed data. The electrode map on the left shows all channels recorded in the original study, with channels used in the model filled in red and channels used to calculate the heartbeat-evoked potential marked with dashed circles.

To further examine whether the model reconstructed physiologically meaningful structure beyond the waveform level, we compared the actual physiological signals from the training datasets with the reconstructed data pooled across both within-cohort and unobserved test sequences. The two datasets showed substantial overlap in several canonical summary properties (Fig. 2c), including the mean power spectral density across EEG channels, the distribution of mean RR interval, and the distribution of within-sequence inhalation-phase duration. These results indicate that the reconstructed data preserved meaningful statistical properties of each modality. Furthermore, heartbeat-evoked potentials, serving as a representative marker of BBI, showed similar waveform profiles between the actual and reconstructed data, with a typical early component clearly evident in the model outputs (Coll et al., 2021). Thus, the generative process reproduced not only the superficial multimodal waveforms but also their underlying physiological architecture, including a critical signature of BBI. These reconstructions highlight the model’s ability to represent multimodal physiological data that combines multiple timescales with both deterministic and stochastic components.

### Multimodal-associative layer latent variables capture multimodal coupling dynamics

Second, building on this generative capability, we evaluated whether the intermediate associative layer captures multimodal coupling. We conducted the ablation study by removing each probabilistic latent variable in turn and evaluating its influence on the generation of each modality (see Methods section). Briefly, we fixed their activities to zero during the generation process and measured how the reconstruction error for each signal was increased using a paired t-test between the full model and ablation model.

Fig. 3 shows the percentage of total latent variables in each layer that actively influence each modality, indicating the extent to which they represent a specific modality. The ablation of latent variables in the executive layer revealed that they play distinct functional roles in representing individual identity and global states. Contrary to the expectation of a uniform global influence, the latent variables were categorized into ‘EEG only’ (one variable), ‘EEG + ECG’ (two variables), and ‘None’ (one variable). The category ‘None’ indicates that ablating the variable did not produce a significant increase in reconstruction error in any modality under the present analysis. This distribution suggests that the task-condition and individual-trait structure self-organized in the executive layer is reflected primarily in brain dynamics and brain–heart interactions, rather than in a uniform influence across all modalities. Furthermore, the absence of top-down influence on respiration implies that basal autonomic rhythms may be represented by lower-level dynamics. However, it is important to note that the lack of significant impact in this analysis does not prove that these variables are completely unrelated to the generation process.

**Fig. 3.**
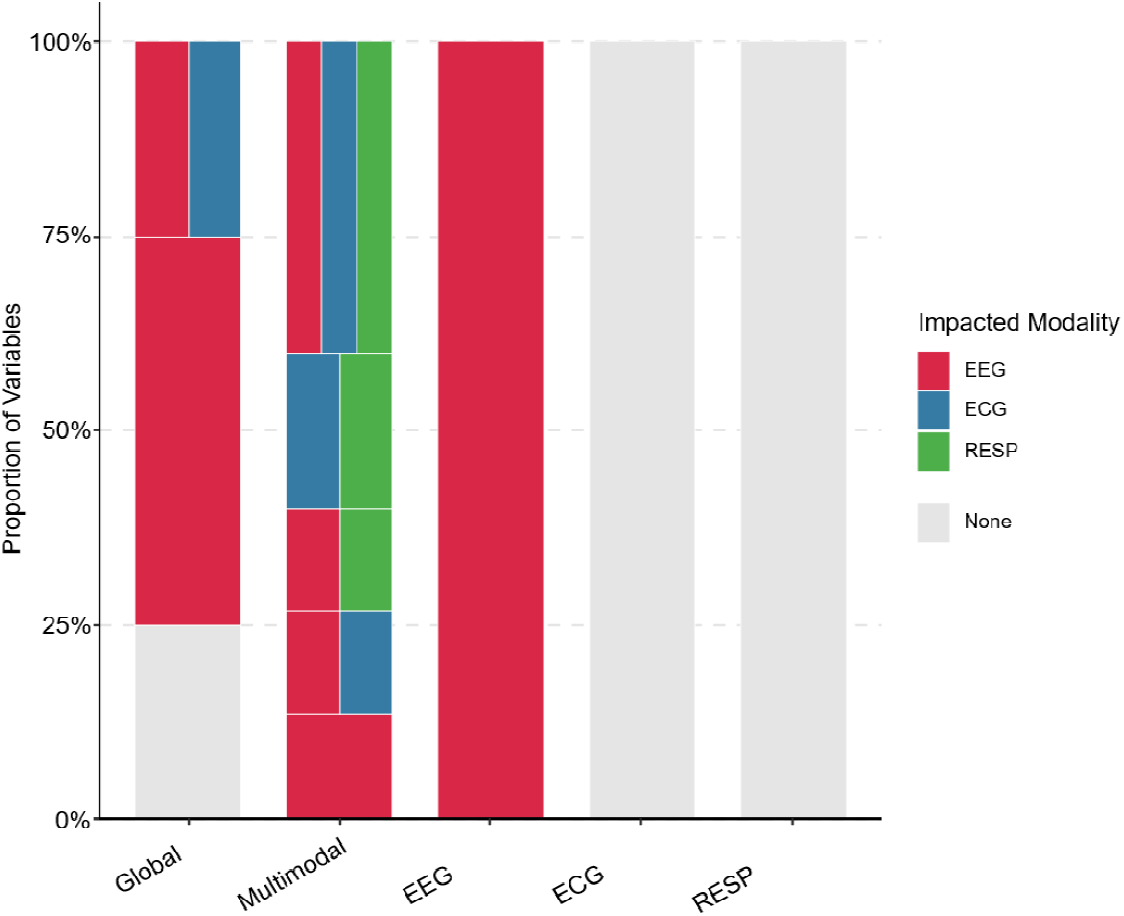
The effects of the latent variables in each layer on the prediction of each modality. Probability of trained latent variables in each module representing either EEG, ECG, and RESP. A simultaneous display of colors means the representation of multiple signals.

Ablating latent variables in the multimodal-associative layer revealed that these variables contributed to the generation of physiological data in three distinct ways. The six variables demonstrated a global integration role, where their ablation simultaneously disrupted the generation of all three modalities. The other eight variables exhibited cross-modal influence, affecting pairwise combinations such as EEG–ECG, EEG–RESP, or ECG–RESP. The remaining single variable impacted only the EEG signal. This dominance of global and cross-modal influences—as opposed to isolated, modality-specific processing—indicates that this intermediate layer is highly specialized for linking brain and bodily signals. Consequently, these results show that the multimodal-associative layer primarily functions to represent the BBI.

Ablation of the probabilistic latent variables in the sensory modality layers had no significant impact on prediction generation, except for the EEG module. However, this result does not imply that these modalities are uninvolved in the data generation process. It is important to note that our ablation targeted only the probabilistic latent variables. In the PV-RNN architecture, predictive outputs are generated from the lowest layer via deterministic variables (*h* and *d*). The internal state *h*_*t*_ at each time step is determined by both top-down inputs (Eq. (1)) and local inputs. Therefore, if higher layers predict the signal dynamics and provide strong top-down signals, the contribution of local probabilistic states becomes negligible, leading to the ablation effect seemingly disappearing. Physiologically, respiration and heart rate are intrinsically linked (Yasuma & Hayano, 2004), and their activities are often reflected within EEG signals (Heck et al., 2016; Schandry et al., 1986; Zaccaro et al., 2024). These results likely reflect the inherent characteristics of these signals: while EEG possesses unique components and complex dynamics not found in other modalities, ECG and respiration exhibit less unique fluctuation and are largely explainable by other modalities or top-down predictions.

### Emergence of multiscale, bidirectional, and variable-lag dynamics in latent associative representations

Furthermore, to demonstrate how the proposed model overcomes the four challenges of BBI—nonlinearity, multiscale dynamics, bidirectionality, and unknown time lags—we examined the dynamics of the probabilistic latent variables in the multimodal-associative layer. We specifically focused on those variables identified as essential for the joint reconstruction of EEG and bodily signals. Visual inspection of their temporal evolution revealed that these variables exhibit complex, state-dependent fluctuations, reflecting the non-linear integration of cross-modal information (Supplementary Fig. 2). Furthermore, spectral analysis demonstrated that these latent variables spontaneously organized into a multiscale temporal structure (Fig. 4a). Some variables mainly reflected slow-frequency dynamics (<1 Hz), likely corresponding to autonomic rhythms such as respiration, while others captured faster fluctuations, thereby enabling the model to manage the diverse timescales inherent in BBI.

**Fig. 4.**
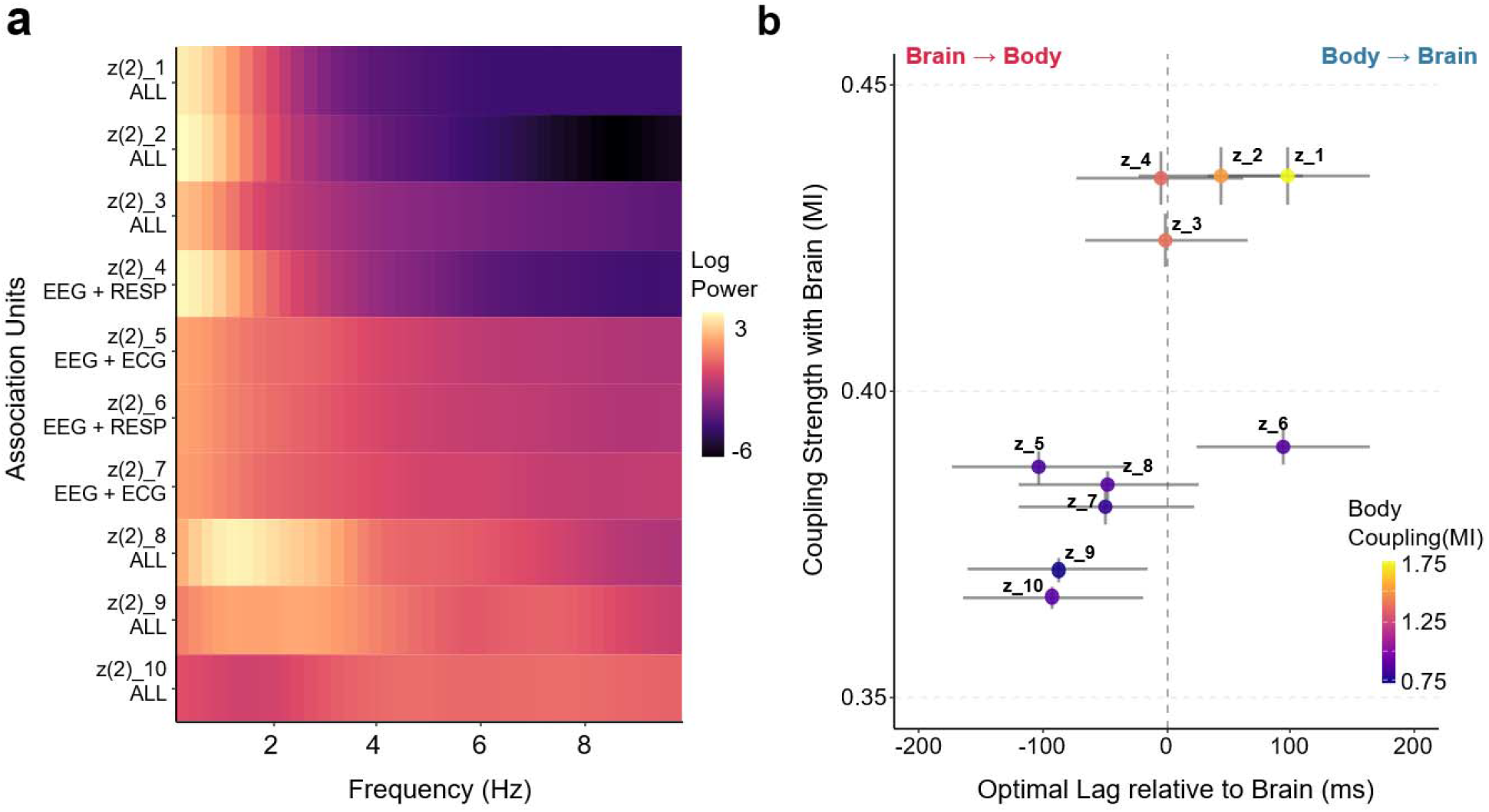
Multiscale dynamics, directional asymmetry, and diverse time lags of latent variables in multimodal-associative layer. **a**. Multiscale spectral properties of the latent variables related to brain and bodily signals in the multimodal-associative layer (z(2)). The power spectral density (PSD) of the 10 variables identified as reflecting the BBI. Variables are sorted from top to bottom based on the ratio of low-frequency power (<1 Hz). **b**. Directionality and coupling strength of interaction. The relationship between the optimal time lag relative to global brain activity and mutual information (MI). The x-axis represents the lag at which MI is maximized. The y-axis indicates the coupling strength with the brain, and the color scale indicates the coupling strength with the body (max MI with ECG or Respiration). Error bars denote standard error (SE).

To determine the directionality of interactions mediated by the multimodal-associative layer, we quantified the Lagged Mutual Information (MI) between each latent variable and global EEG activity, summarized as Global Field Power (GFP). By varying the time lag, we identified the optimal temporal offset that maximized information sharing (Fig. 4b). Negative optimal lags indicated that brain activity preceded changes in the latent variable. Because the ablation analysis showed that associative-layer variables are necessary for the joint generation of brain and bodily signals (Fig. 3), these brain-leading variables can be interpreted, with caution, as reflecting an efferent pathway from the brain to the associative layer to the body. In contrast, positive optimal lags indicated variables whose activity preceded global EEG fluctuations. As these same variables were also coupled to somatic signals, they may reflect afferent pathways from the Body to the Associative to the Brain. Although these pathway labels remain inferential, the broad distribution of optimal lags indicates that the model captured variable temporal delays characteristic of physiological loops. Together, these findings suggest that the associative layer acquired a nonlinear, multiscale, and bidirectional representation of BBI.

### Attentional precision enhancement of brain-body interactions aligns with psychological and psychiatric traits

Third, we evaluated the construct validity of the extracted latent states by assessing their correspondence with the experimentally manipulated attentional conditions. To demonstrate that the proposed model yields an interpretable and theory-consistent index of BBI, we showed these dynamic fluctuations (posterior mean and standard deviation) time-locked to the heartbeat R-wave, providing a clear intuitive representation of brain–body coupling (Fig. 5a). In predictive-coding accounts of perception, directing attention to a target is expected to increase the precision of the corresponding sensory signals (Friston, 2010; Friston & Kiebel, 2009). BF required participants to allocate attention to bodily signals, whereas SF directed attention to external auditory input. If the associative latent variables identified in the PV-RNN as being involved in joint brain-body reconstruction genuinely capture BBI-related structure, their condition dependence should be consistent with this conceptual manipulation. We therefore asked whether interoceptive attention (BF) would be reflected in the trained PV-RNN as reduced posterior uncertainty in the multimodal-associative layer compared with exteroceptive attention (SF). We focused on the state-level modulation of uncertainty by averaging the posterior standard deviation (*σ*) across all time steps within each sequence. Furthermore, we restricted our analysis to the variables identified as essential for the joint reconstruction of EEG and bodily signals, same with previous section. Consistent with the predictive-coding account, the average uncertainty of latent variables reflected BBI was significantly lower during the BF condition than the SF condition (Fig. 5b; *t*(32) = 2.18, *p* = 0.037), indicating a sustained enhancement of attentional precision at the multimodal-associative layer during interoceptive focus.

**Fig. 5.**
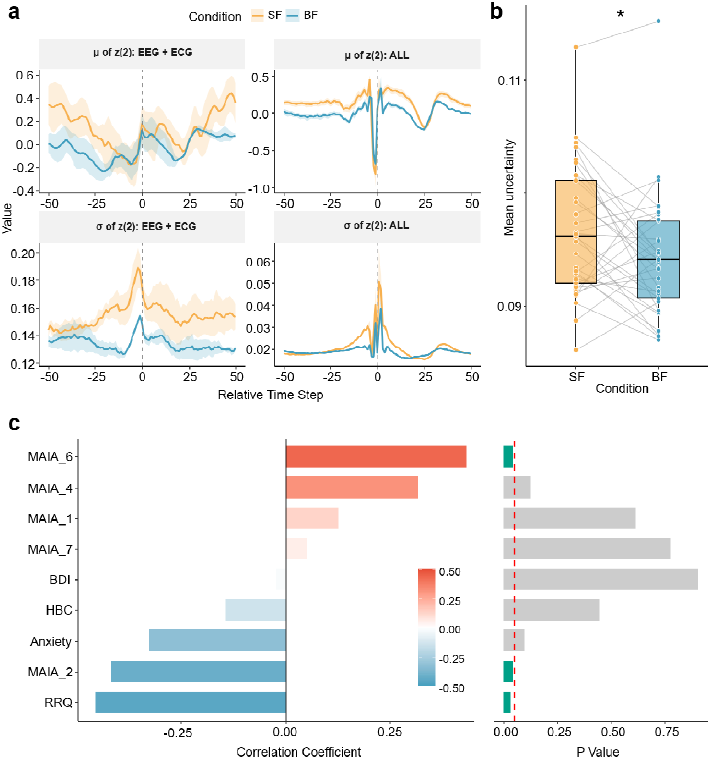
Interoceptive focus enhances association-layer precision and relates to psychological and psychiatric traits. **a**. Example of a time series of posterior uncertainty in the multimodal-associative layer. b. Multimodal-associative layer posterior uncertainty (mean σ of each latent variable) is reduced in interoceptive attention compared to exteroceptive attention. **b**. Correlations between primary principal component of precision enhancement ( ) and psychological/ psychiatric measures. HBC: heartbeat counting task. RRQ: Rumination-Reflection. Questionnaire.

Next, to assess the external validity of this model-derived index, we evaluated whether individual differences in BF-induced precision enhancement relate to psychological and clinical traits theoretically linked to interoceptive processing (Mehling et al., 2012; Paulus & Stein, 2010). For each participant, we computed an uncertainty modulation score as Δ*σ* = *σ*_*SF*_ − *σ*_*BF*_, such that larger positive values indicate stronger uncertainty suppression during BF. To ensure statistical robustness given our relatively small sample size (*N* = 33), we calculated Pearson’s correlation coefficients and assessed their significance using non-parametric permutation testing with 10,000 iterations, and corrected p-values using False Discovery Rate (FDR) correction within each trait category to control for multiple comparisons. Our analysis revealed significant associations in both directions (Fig. 5c; Supplementary Table1). Specifically, greater precision enhancement was significantly associated with lower rumination (RRQ; *r* = - 0.45, *p* = 0.033) and a lower tendency to distract oneself from uncomfortable sensations (MAIA Not-Distracting; *r* = -0.41, *p* = 0.046). Conversely, a significant positive association was observed with adaptive interoceptive regulation (MAIA Self-Regulation; *r* = 0.43, *p* = 0.046). Furthermore, trait anxiety showed a marginally significant negative trend (STAI-Trait; *r* = -0.32, *p* = 0.097), whereas depressive symptoms (BDI) and the other MAIA subscales did not show significant relationships. Together, these results indicate that the multimodal association-layer posterior uncertainty provides a condition-sensitive, interpretable representation of BBI, whose individual differences robustly align with clinically and theoretically relevant traits.

### Higher-level latent representation dissociates attentional condition and individual differences

Finally, we investigated whether the highest-level latent representations spontaneously segregate global sequence-level information, capturing both within-subject attentional conditions (BF or SF) and between-subject individual differences in a fully data-driven manner. We visualized the latent space using a two-dimensional t-SNE embedding (Fig. 6a,b), where each point corresponds to one sequence. Notably, although these labels were not used for model training, the latent space exhibited structured organization that aligned with externally defined sequence labels (condition and individual): coloring points by attentional condition or, separately, by participant identity revealed label-consistent grouping patterns. To quantify this organization, we computed Silhouette scores in the original global state latent variables and assessed significance against permutation-based null distributions obtained by randomly shuffling labels. Although the absolute silhouette values were low, the observed scores were significantly higher than chance for both condition and participant identity (Fig. 6c,d; condition: p < .001; individual identity: p <.001), indicating that the global state layer latent variables form a self-organized representation that reflects both state-like (condition) and trait-like (individual) factors.

**Fig. 6.**
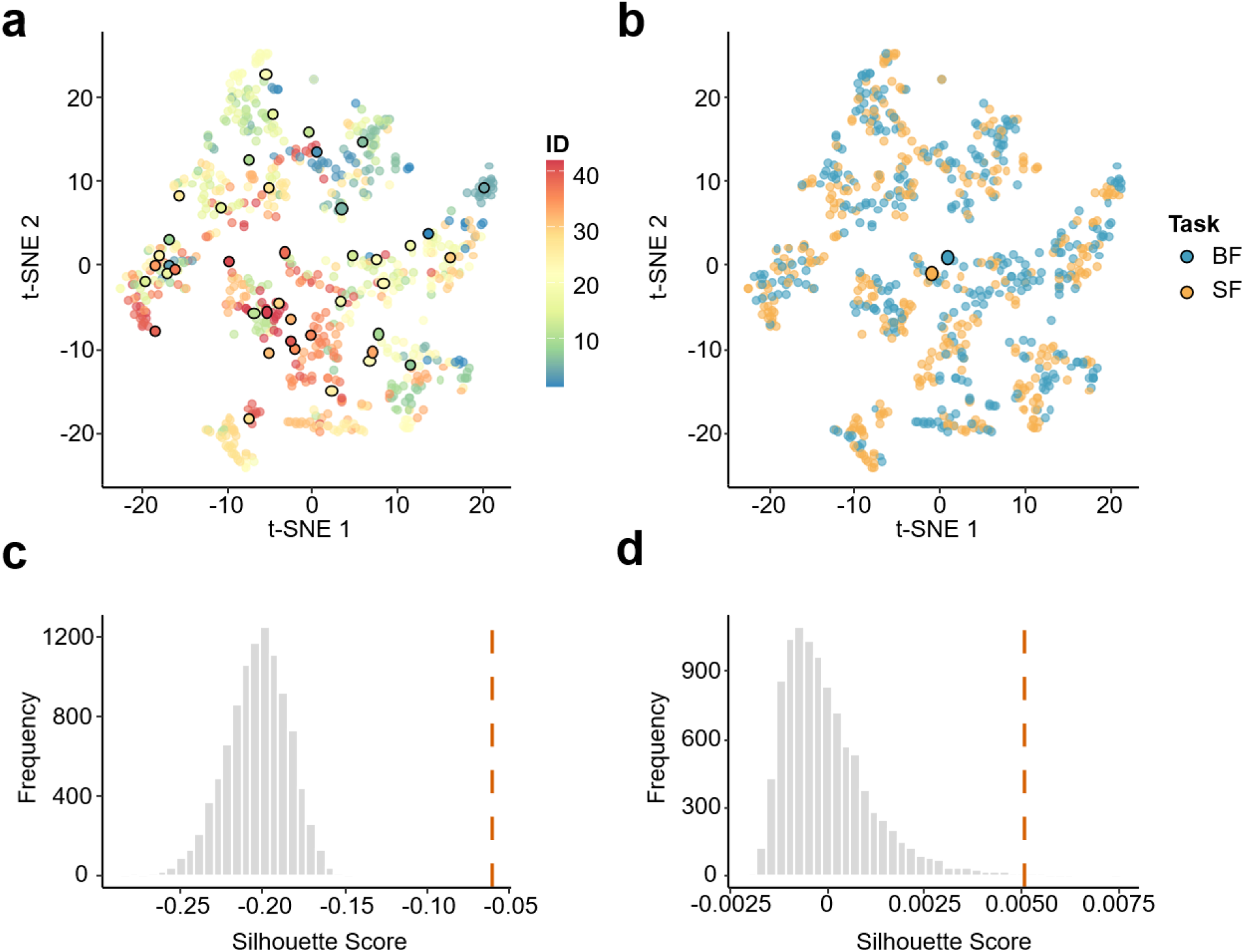
Global state latent variables self-organize to reflect attentional condition and individual identity. **a**. Two-dimensional t-SNE embedding of global state layer latent variables. Each point corresponds to one sequence and is colored by participants ID. Larger points with black outlines indicate the centroid of each participant’s cluster. **b**. The same embedding as in Fig.3a, with points colored by attentional state. Larger points with black outlines represent the centroid of each attentional condition. **c**. Silhouette scores computed in the original global state latent space for condition and identity, compared with permutation-based null distributions obtained by random labels. Vertical dotted lines indicate observed values of Silhouette score.

## Discussion

In this study, we established a data-driven and interpretable framework for modeling BBI during interoceptive and exteroceptive attention using a PV-RNN trained on simultaneous EEG, ECG, and respiratory signals. First, the model reliably reconstructed multimodal physiological time series not only for held-out sequences within the training cohort but also for previously unobserved participants, indicating that the inferred latent representations were not driven by sequence-specific overfitting. Second, analyses of the intermediate multimodal-associative layer showed that the model captured cross-modal coupling dynamics linking brain and bodily signals. These dynamics, which reflect brain and bodily signals, successfully captured the inherent characteristics of BBI, including nonlinearity, multiscale dynamics, bidirectionality, and unknown lags. Third, posterior uncertainty in this associative layer decreased during interoceptive focus, and individual differences in this precision modulation were associated with subjective interoceptive regulation and psychiatric traits. Finally, the highest-level latent representation self-organized to reflect both attentional condition and individual differences. Together, these findings support PV-RNN as a unified generative framework for extracting BBI as hierarchical latent dynamics from continuous multimodal physiology.

The validity of the model extends beyond mere point-by-point waveform fitting. It successfully captured multimodal physiological dynamics with fundamentally distinct temporal properties, including fast irregular EEG fluctuations, rhythmic respiratory activity, and ECG waveforms characterized by relatively fixed shapes but variable timing (Fig. 2a). This reconstruction was not superficial, as the model accurately reproduced the underlying statistical characteristics of each modality (Fig. 2c) and even replicated canonical indices of BBI, such as heartbeat-evoked potentials (Fig. 2d). By employing a generative approach, we represented these diverse physiological processes within a single unified framework, without reducing them to discrete events, manually defined features, or pairwise summary statistics. The successful reconstruction of these signals implies that the latent space preserved sufficient underlying structure to intrinsically reproduce the continuous multimodal stream. Furthermore, the probabilistic nature of the latent representations likely contributed to this robust performance. Because the model can treat a portion of the inherent noise and high variability in physiological signals as uncertainty, it effectively avoids discarding essential generative information, thereby enabling the extraction of robust BBI-related structures from highly complex and noisy multimodal data.

The most important mechanistic result is that the multimodal-associative layer appears to function as the representation of BBI. Our ablation study revealed that two-thirds of the latent variables in the multimodal-associative layer contributed to reconstructing both brain and bodily signals. Analyses of activities of these variables, identified as reflecting BBI, demonstrated their nonlinearity (Supplementary Fig. 1) and distinct oscillation peaks (Fig. 5a). Moreover, the bidirectional influences and temporal lags related to brain activity were estimated in a data-driven manner (Fig. 5b). The successful acquisition of these complex BBI dynamics can be attributed to two specific architectural properties of the PV-RNN. First, the intrinsic recurrent and state-dependent nature of the RNN enables the latent space to approximate nonlinearities that linear models fail to capture. Its memory mechanism bridges variable time lags between the brain and body, while its dynamic internal states allow the model to represent complex, context-modulated interactions. Second, the spatiotemporal hierarchy plays a crucial role in integration. The emergence of distinct oscillation peaks within the multimodal-associative layer indicates that distinct timescales from the lower modality-specific layers are integrated into a single latent dynamic. This layer functions as a common source for top-down predictions. By minimizing prediction errors through predicting the future states of both brain and body from their shared past states, this hierarchical constraint effectively captures the bidirectional causality and variable lags inherent in BBI dynamics.

The observed enhancement in BBI precision during interoceptive focus likely stems from multifaceted mechanisms. Since latent variables in the multimodal-associative layer capture brain and bodily activities, this modulation arises not from isolated peripheral or cortical events, but fundamentally from their bidirectional interplay. First, regarding physiological modulation, respiratory sinus arrhythmia—where heart rate accelerates during inspiration and decelerates during expiration—represents a fundamental coupling between cardiac and respiratory systems (Yasuma & Hayano, 2004). This coupling is known to strengthen when respiration becomes slower and more regular (Hirsch & Bishop, 1981), a pattern often induced by voluntary attention to breathing (Wielgosz et al., 2016). Consistent with this, our data showed a reduction in respiratory rate during the focus on their respiratory compared to auditory stimuli (Shinagawa et al., 2025). Such stabilization of cardiorespiratory coupling likely enhances the predictability of these signals and the precision of their neural representation. Second, the modulation of cortical processing via attentional allocation plays a critical role. Previous studies suggest that interoceptive attention alters peripheral physiology and the gain and precision of the cortical processing of interoceptive signals (Petzschner et al., 2019). Specifically, the heartbeat-evoked potential, a cortical response to cardiac afferents, is modulated by interoceptive attention and is reported to depend on the respiratory phase (Zaccaro et al., 2024, 2022). This suggests that top-down attention enhances the respiratory-phase-dependent gating of cardiac representation in the brain. Consequently, such bidirectional interplay between the regularized cardiorespiratory signals and the modulated cortical processing may underlie the stabilized latent inference of brain and bodily activities within the multimodal-associative layer.

Individual differences in BBI precision enhancement were correlated with subjective adaptive control abilities measured by a questionnaire. We observed positive correlations with the Self-Regulation subscales of the MAIA (Mehling et al., 2012), which measure the ability to regulate distress by attending to bodily sensations. Since this measure reflects the subjective capacity to voluntarily direct attention toward the body, they align with our finding that participants who effectively focused on respiration exhibited greater precision enhancement. Conversely, a negative correlation was observed with the Not-Distracting subscale. This scale reflects the tendency not to ignore or distract oneself from sensations of pain or discomfort (e.g., a low score on the item “I distract myself from sensations of discomfort” results in a high score). Conceptually, high scores on this scale indicate a susceptibility to being passively captured by salient, bottom-up interoceptive signals, rather than exercising top-down attentional control. Since the experimental task required the active, top-down regulation of attention—conceptually the opposite of being passively captured by bottom-up sensations—these opposing processing dynamics likely explain the observed negative correlation. Together, these results highlight that the model-derived BBI precision effectively captures the individual capacity for the top-down, active precision control of interoceptive inference.

Crucially, this interpretation of precision enhancement provides a coherent framework for understanding the observed negative correlations with psychiatric vulnerabilities, specifically rumination and the trend with trait anxiety. Predictive coding framework conceptualizes altered processing of bodily signals in anxiety and depression— conditions having rumination as a core component of symptoms—as aberrant precision weighting, wherein a failure in precision modulation amplifies interoceptive prediction errors and impedes flexible adaptation to noisy ascending signals (Barrett & Simmons, 2015; Paulus & Stein, 2010). While theoretical accounts posit altered BBI in these conditions, empirical evidence relying on conventional behavioral indices of cardiac interoception (e.g., heartbeat counting task) has remained inconsistent (Eggart et al., 2019; Jenkinson et al., 2024). This inconsistency likely stems from the inherent limitations of static behavioral tasks, which capture generalized sensory accuracy and suffer from low construct validity. Overcoming these limitations, our generative modeling approach isolates the top-down control of BBI by extracting task-dependent precision modulation from multimodal physiological data, allowing us to translate the abstract concept of aberrant precision into a quantifiable dynamic. Specifically, individuals prone to rumination demonstrated a marked difficulty in enhancing BBI precision during interoceptive attention. This finding suggests that the altered processing of bodily signals in psychiatric conditions is not simply a static sensory deficit or constant hypersensitivity, but fundamentally a failure of context-dependent precision reallocation (Stephan et al., 2016). Adaptive regulation requires flexibly modulating precision weighting across internal and external signals according to task demands. The inability of highly ruminative individuals to make this shift indicates that their inferential system is too rigid to redistribute precision across changing contexts. For instance, when confronted with uncomfortable physical sensations or distress, individuals with such computational rigidity may fail to adaptively down-regulate their bodily signals, becoming trapped in the unpleasant state rather than flexibly shifting their interoceptive focus. This computational deficit thus provides a mechanistic explanation for the physical and emotional inflexibility commonly observed in these conditions, suggesting that such psychiatric vulnerabilities fundamentally emerge from a systemic lack of dynamic flexibility in the BBI.

It is necessary to address a methodological concern in multimodal modeling, whether the shared latent space captures common noise sources, such as cardiac or respiratory motion artifacts contaminating the EEG signal (Urigüen & Garcia-Zapirain, 2015). At first, we implemented a preprocessing pipeline—including 0.5-Hz high-pass filtering to remove low-frequency drifts and independent component analysis to eliminate cardiac artifacts—which substantially attenuates the influence of these non-neural components within the EEG signals. Beyond these signal-level corrections, the empirical relationships detailed above provide the strongest conceptual evidence against the artifact hypothesis. Within our model’s hierarchical architecture, latent dynamics in the associative layer are governed by the higher-level global state layer, which self-organizes to encode individual identity and attentional context (Fig. 3). This top-down control suggests that the extracted dynamics reflect intrinsic, context-dependent representations rather than sequence-specific noise. Furthermore, the magnitude of the precision modulation systematically correlated with subjective adaptive body control and psychiatric traits. Residual physiological noise cannot account for such coherent dependencies on hierarchical cognitive contexts and multifaceted clinical phenotypes. Therefore, these findings demonstrate that our model captures conceptually valid BBI dynamics rather than physiological signal artifacts.

In this study, we developed a PV-RNN to extract complex interactions among EEG, ECG, and respiration. While our current focus was on reconstructing and interpreting these BBI dynamics, the model’s inherent generative capacity provides a highly manipulable platform for in silico physiological simulations. Crucially, as a foundational proof of this potential, our analysis revealed that the highest-level latent variables spontaneously self-organized to reflect both attentional conditions and individual differences without explicit labels. By naturally segregating broader context at the highest level while preserving moment-to-moment generative BBI mechanisms at lower levels, our framework achieves robust state discrimination alongside mechanistic interpretability. This structural capability makes it uniquely suited for in silico experiments. Existing methods to manipulate visceral activity typically rely on exogenous stimulation (Harrison et al., 2021; Smith et al., 2021); in contrast, our framework can simulate spontaneous autonomic regulation simply by modulating latent variables and generating the corresponding data. For instance, by artificially inducing specific physiological events—such as transient cardiac acceleration—within the latent space during resting or task states, researchers could systematically investigate how these events influence spontaneous brain activity or cognitive processing. This in silico approach effectively circumvents the ethical and practical barriers associated with direct human interventions. Extending beyond fundamental cognitive research, this framework has potential for clinical applications. By training on data from individuals with specific conditions—such as depression or anxiety, where interoceptive processing is altered—it enables the creation of patient-specific models for virtual therapeutic simulations. Capable of replicating individual- and state-specific dynamics, broadening the horizon of BBI research from elucidating basic cognitive mechanisms to realizing high-precision, personalized medicine in psychiatric disorders.

In recent years, the framework of predictive coding has increasingly conceptualized interoception, viewing the perception and regulation of bodily states as inference (Seth et al., 2011). While computational studies using behavioral data have suggested that the estimation of interoceptive precision is altered in psychiatric conditions (Smith et al., 2020), empirical validation of these mechanisms using continuous physiological data has remained limited. For example, although phenomena such as the modulation of HEP amplitude by interoceptive attention have led to the proposal that the HEP reflects precision-weighted prediction errors for individual heartbeats, this hypothesis has not yet been fully established (Ainley et al., 2016; Petzschner et al., 2019). Overcoming these limitations, the PV-RNN inherently incorporates continuous predictions and their variance, providing a unique advantage for operationalizing these theoretical constructs directly from multimodal data. As demonstrated by our analysis of psychiatric traits— which serves as a compelling proof-of-concept—the model successfully translates abstract computational mechanisms into measurable neural dynamics. Crucially, the scope of this generative framework extends far beyond psychopathology. It offers a foundational tool for empirically testing a broad spectrum of cognitive functions and autonomic regulations hypothesized by predictive coding. Furthermore, it is also expected to extend this approach not only to verify theoretical predictions but also to experimentally explore multimodal agent models incorporating the bodily state, based on the predictive-coding framework (Idei et al., 2025).

In summary, we demonstrated that a hierarchical generative model of multimodal physiological data extracts latent dynamics that capture the core features of BBI, confirming their conceptual validity. This framework is highly flexible, providing a foundation to include other physiological or behavioral signals in the future. Because the model is inspired by the predictive-coding framework, it is consistent with current computational theories of the brain and helps us test existing ideas about cognitive function and autonomic control. Additionally, by isolating specific neural signatures linked to bodily rhythms, this approach directly addresses traditional psychophysiological interests, such as extracting the EEG components of spontaneous respiration. Furthermore, its ability to generate data creates new opportunities for in silico experiments, hypothesis generation, and clinical application. Therefore, this approach provides a powerful tool to expand the focus of cognitive neuroscience— which has long centered on the brain—to fully include the dynamic relationship between the brain and the body.

## Methods

These data were originally described in (Shinagawa et al., 2025), where full experimental details can be found. The dataset includes the simultaneous EEG, ECG, and respiratory recordings acquired during exteroceptive (SF) and interoceptive (BF) attention tasks. In SF, participants responded as quickly as possible to randomly presented pure tones (inter-tone interval: 1.2–2.0 s), whereas in BF they responded at the end of each exhalation; the same pure tones were also presented in BF to equate the auditory environment. In both conditions, participants performed self-reports when they noticed their attention had drifted away. We extracted the periods 1–5 s preceding these reports, representing intervals when participants were presumably focused on the task (i.e., on sounds or their breath).

The original EEG data were recorded from 64 electrodes distributed across the entire scalp. To balance computational efficiency with spatial coverage, we selected 19 representative electrodes for subsequent analysis. These electrodes were chosen to ensure comprehensive cortical coverage, incorporating primary sites within the frontal, temporal, parietal, and occipital lobes. In contrast, all available ECG and respiratory channels were utilized to fully capture autonomic dynamics. While the primary signals were acquired at 500 Hz, they were downsampled to 100 Hz to reduce the computational load for the recurrent neural network training. Each resulting sequence consisted of 400 time points (4 s duration), comprising 19 EEG, 2 respiratory, and 1 ECG channel. Continuous EEG signals were filtered using a 0.5–50 Hz bandpass filter. Power line noise at 50 Hz was removed, and artifact correction was performed using Independent Component Analysis (ICA) to exclude components with an 80% or greater probability of being noise (e.g., muscle activity, cardiac artifacts). ECG data were preprocessed with a 3–30 Hz bandpass filter to facilitate R-peak detection. Respiratory signals were mean-smoothed and detrended to determine respiratory phases. Detailed preprocess pipeline is in (Shinagawa et al., 2025)).

While the original dataset included 44 participants, we applied a inclusion criterion to ensure the robustness of the staged evaluation (within-cohort and unobserved-participant tests) for evaluating generalization performance across novel, previously unobserved individuals. For the training cohort, we included 33 participants who each provided at least 13 valid mind-wandering (MW) reports for both the SF and BF conditions. From each participant’s data, 10 sequences per condition were assigned to the training set, while 3 sequences per condition were reserved for validation. For the unobserved-participant test, we selected 9 additional participants from the remaining pool who met the criterion of providing at least 3 reports per condition.

### Model Architectures

In this section, we describe the mathematical details of top-down prediction generation and bottom-up parameter updates using the PV-RNN (Fig. S1).

#### Top-down prediction generation process

Prediction generation was performed in a top-down manner using a network hierarchy. The internal state *h*_*t,i*_ and output *d*_*t,i*_ of the *i*th deterministic variable (recurrent unit) in 6 each network module at time step *t* (*t* ≥ 1) is calculated as:

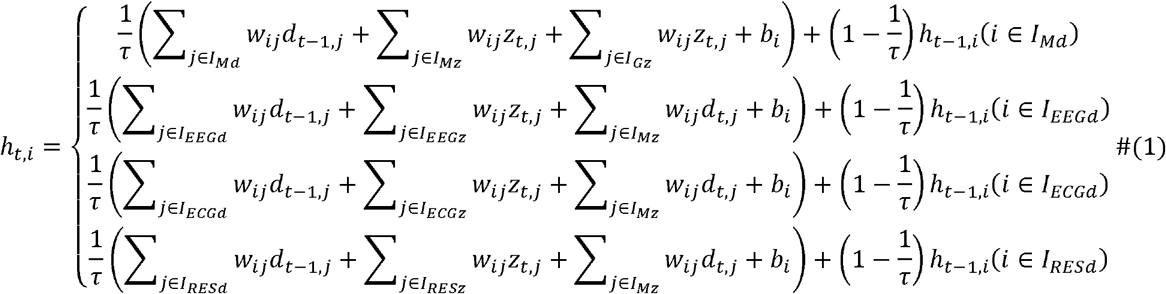

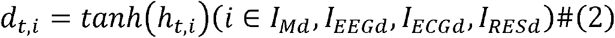

where *I*_*Md*_,*I*_*EEGd*_,*I*_*ECGd*_,*I*_*RESd*_ and Ird, are the index sets of deterministic variables in the multimodal-associative module, EEG module, ECG module, and RESP module, respectively. *I*_*G*z_, *I*_*Mz*_, *I*_*EEGz*_, *I*_*ECGz*_,*I*_*RESz*_ are the index sets of probabilistic latent states in the global state module, multimodal-associative module, EEG module, ECG module, and RESP module, respectively. *w*_*ij*_ is the weight of the synaptic connection from the *j*th neuron to the *i*th neuron; *z*_*t,j*_ is the output of *j*th latent states at time step *t*;*τ* is the time constant of the deterministic variable; and *b*_*i*_ is the bias of the *i*th deterministic variable. A deterministic variable with a small time constant *τ* has a tendency to change its activity rapidly, while that with a large time constant has a tendency to change its activity slowly (Yamashita and Tani, 2008). We set the initial internal states of the deterministic variables *h*_0,*i*_ (*i* ∈ *I*_*Md*_, *I*_*EEGd*_, *I*_*ECGd*_, *I*_*RESd*_) to 0 (*d*_0,*i*_ is also 0).

The probabilistic latent state *z*_*t*_ in each module is assumed to follow a multivariate Gaussian distribution with a diagonal covariance matrix, meaning *z*_*t,i*_ and *z*_*t,i*_ are independent (*i,j* ∈ *I*_*Gz*,_ *I*_*Mz*_ *I*_*EEGz*_ *I*_*ECGz*_ *I*_RESz_ ∧*I* ≠ *j*). Here, the mean *μ*_*p,t,i*_ and standard deviation *σ*_*p,t,i*_ of the prior distribution *p*(*z*_*t,i*_) in all modules are calculated as below:

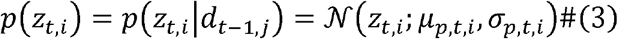

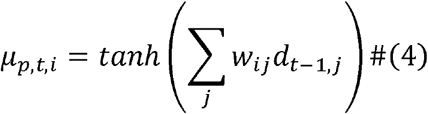

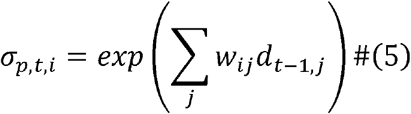

Here, the prior distribution in each module is calculated exclusively from the previous deterministic states within the same module. The posterior distribution in each module is calculated as follows:

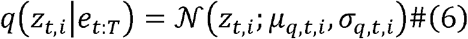

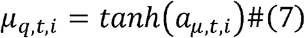

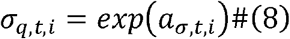

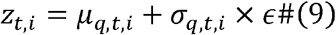

Here, *T* represents the last time step of the time window. *a*_*t*_ is the adaptive internal state of the latent variables representing the posterior distributions. The adaptive variables *a*_*t*_ are determined by error signals *e*_*t:T*_ propagated using the back propagation through time (BPTT) algorithm (Friston & Kiebel, 2009). The adaptive variables *a*_*t*_ are initialized by the corresponding initial internal states of the latent variables that represent the prior distributions. In Eq(9), by sampling *ϵ* epsilon from *𝒩* (0,1), the latent state *z*_*t,i*_ is obtained.

Finally, predictions for EEG, ECG, and RESP signals were individually generated from the EEG, ECG, and RESP modules, respectively as follows:

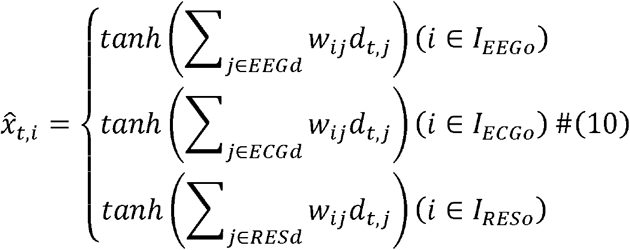

Here, *I*_*EEGo*_, *I*_*ECGo*_, *I*_*RESo*_, are the index sets of the output units for the EEG, ECG, and RESP predictions, respectively.

#### Parameter updates through bottom-up optimization

Optimization processes in both training and test phases have been implemented by employing the negative Evidence Lower Bound (ELBO) as the loss function. This negative ELBO is equivalent to the variational free energy (Friston & Kiebel, 2009) in the context of brain computation theory, and is hereafter identified as *F*_*t*_. In the framework of PV-RNN, *F*_*t*_ is derived as follows (Idei et al., 2022; Takahashi et al., 2025).

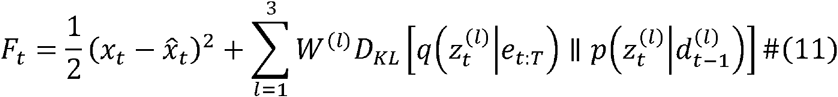

*l* denotes the hierarchical level from bottom to top (i.e. sensory to global state). The first term of this equation represents the prediction error, while the second term denotes the influence of the prior 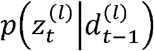 on the update of the posterior 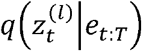 of the latent state. Consequently, in an optimization process aimed at minimizing *F*_*t*_, the update of the posterior progresses in a manner that minimizes the prediction error under the influence of the prior. The hyperparameter *W*^(*l*)^, referred to as the meta-prior (Ahmadi & Tani, 2019), serves to balance the prediction error and the divergence between the prior and posterior, and can be individually set for each hierarchical level. In this study, we set the meta-prior to a uniform value across all levels (*W*^(1)^ = *W*^(2)^ = *W*^(3)^ = 0.001). If the meta-prior is too small, overfitting occurs, whereas if the meta-prior is too large, the reconstruction error does not decrease sufficiently (Ahmadi & Tani, 2019). This optimization by minimizing *F*_*t*_ was carried out during both the training and test phases.

During the training phase, synaptic weight *w* and the adaptive variables *a*_1:*T*_ are updated to minimize the free energy 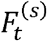 over all time steps and target sequences as follows. On the other hand, during the data assimilation phase, only the posterior is updated, while the weights remain fixed.

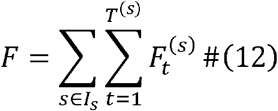

#### Hyper-parameter setting

The dimensions of the latent variables in the EEG, ECG, RESP, and multimodal-associative modules were, *N*_*EEG*_ = 15, *N*_*ECG*_ = 2, *N*_*RESP*_ = 2, *N*_*multi*_ = 15 respectively. These numbers were determined by a preliminary experiment in which we increased the dimension of the latent variable in the multimodal-associative module individually and searched for the minimal setting for successful reconstruction of the learning and test data. Regarding the global state layer, the dimension was set to *N*_*Global*_ = 4. This choice was grounded in an iterative evaluation where the dimensionality was varied from 2 to 5 (Supplementary Fig. 3). We assessed the capacity of the global layer space to encode task-relevant information—specifically participant identity and attentional state—using the Silhouette score. *N*_*Global*_ = 4 was selected as the minimal configuration that sufficiently captured these hierarchical structures without overfitting, allowing the model to represent both stable individual traits and state-dependent attentional contexts within its highest latent level. The number of deterministic recurrent variables in the EEG, ECG, RESP, and multimodal-associative modules were *N*_*EEG*_ = 40, *N*_*ECG*_ = 20, *N*_*RESP*_ = 20, *N*_*multi*_ = 50 required for reconstructing the learning data.

We incorporated a temporal hierarchy into the model by assigning multiple timescale properties to the deterministic recurrent variables. Specifically, the time constants for the modality-specific modules were set to 2 or 4, while those in the multimodal-associative module were set to 4 or 8. To implement these multiple timescales within each level, the deterministic variables were divided into two equal groups, with each group being assigned one of the two specified time constants. As such, the higher-level modules have a larger time constant (slower neural dynamics) than the lower-level modules have, thus implementing a temporal hierarchy. The synaptic weights were initialized with random values based on the Xavier method (Glorot & Bengio, 2010). The biases of the deterministic recurrent variables were initialized with and fixed to random values following a Gaussian distribution N(0,10), based on a previous study (Idei et al., 2025).

During training, the weights *w* and posterior of latent state values were updated 200,000 times. In the test phase, we updated only the posterior for the latent variables and repeated 20,000 times. Hyperparameters of the Rectified Adam optimizer during learning and test stages were set to their default values: *α*□ = □0.001 (learning rate), *β*_1_□ = □ 0.9, and *β*_*2*_□= □0.999.

### Statistical analysis

#### Clustering methods

We quantitatively assessed how the latent states at the global state layer self-organized into structures reflecting sequence-specific information. To evaluate the degree of this self-organized cluster, we calculated Silhouette scores for each sequence in the original high-dimensional latent space of the global state layer (Fig. 3) (Rousseeuw, 1987). The Silhouette score *S*_*i*_ for a sequence is *i* defined as:

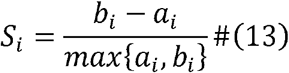

where *a*_*i*_ is the average distance between sequence *i* and all other sequences within the same cluster (e.g., the same participant or condition), and *b*_*i*_ is the minimum average distance from sequence *i* to sequences in a different cluster. *a*_*i*_ and *b*_*i*_ were defined as follows:

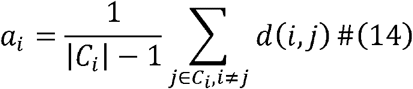

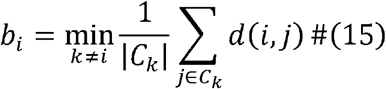

Here, |*C*_*i*_| is the number of points belonging to cluster *C*_*i*_, and *d*(*i,j*) is the distance between data point *i* and *j*. Lower *a*_*i*_ value represents a greater cohesion in cluster, and higher *b*_*i*_ represent a greater separation from adjacent clusters. We performed this analysis across two distinct labeling criteria: (i) Attentional condition to assess state-dependent clustering, and (ii) Participant identity to assess trait-dependent clustering. To statistically validate the observed self-organization against chance, we employed a permutation test. Specifically, a null distribution of Silhouette scores was generated by randomly shuffling the labels across all sequences 10,000 times. The empirical p value was then determined as the proportion of the null distribution exceeding the observed mean Silhouette score.

#### Analysis of Latent Representation

An analysis of the organized latent representation via the ablation of latent variables was conducted (Fig. 4). First, to establish the baseline variability, we used the trained network to generate 20 simulated time series for each signal under different random samplings of the latent states. We then calculated the deviations between these generated time series and the target data. The distribution of prediction error from the 20 iterations for each modality (EEG, ECG, and respiratory) was defined as the baseline difference (BD). Next, to evaluate the impact of individual latent variables, we ablated one variable at a time. Using this lesioned network, we generated 20 simulated time series via random sampling of the remaining latent states. We then calculated the prediction error between these generated time series and the target data. The distribution of prediction error from the 20 iterations for each modality (EEG, ECG, and RESP) was defined as the ablation effect (AE). We compared the distribution of BD and AE using paired t-test. If the AE for a certain modality (EEG, ECG, and RESP) was larger than the corresponding BD, we considered that the ablated latent variable represented the affected modality information. For example, if the ablation of a latent variable of the multimodal-associative module affects EEG and ECG generation, the ablated latent variable is considered to represent both EEG and ECG and is a bimodal latent variable. Finally, we categorized the types of latent variables based on the information represented in each network module and calculated the probability for each type of latent variable (Fig. 4).

#### Analysis of Latent Dynamics and Directionality

To characterize the functional properties of the associative layer, we analyzed the time series of the probabilistic latent variables extracted from the trained model. We computed the Power Spectral Density (PSD) of each latent variable using Welch’s method. Variables were sorted based on the ratio of low-frequency power (<1 Hz) to total power to visualize the hierarchical organization of temporal scales (Fig. 5a). We employed Lagged Mutual Information (MI) to determine the directionality and temporal delay of interactions between the associative variables and brain signals. To avoid the combinatorial explosion of evaluating all channel-latent pairs across multiple time lags, and to capture macroscopic brain state dynamics, we calculated the EEG signal *e*_*t*_ as the spatial standard deviation across all channels, which is equivalent to GEP. For each latent 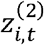 and the EEG signals *e*_*t*_, we computed the MI at various time lags *τ* (ranging from -500 to +500 ms): 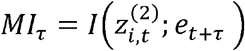, where I(·;·) denotes the Shannon mutual information. The optimal lag *τ*_*opt*_ was defined as the lag maximizing *MI*. A negative *τ*_*opt*_ indicates that brain activity precedes activity of latent variables (top-down), while a positive *τ*_*opt*_ indicates that activity precedes brain activity (bottom-up). The magnitude of the peak *MI* represented the coupling strength with the brain. Similarly, we calculated the *MI* between each unit and somatic signals (ECG and RESP) across lags and indicated by the color scale in Fig. 5b. All *MI* calculations were performed using discrete binning (8 bins, equal-frequency method) to capture non-linear dependencies.

#### Analysis of multimodal association precision

To evaluate the condition-dependent modulation of interoceptive precision, we focused on the posterior uncertainty (*σ*) of the latent variables in the Multimodal-associative layer. We selected a subset of 10 latent variables that were independently identified as capturing BBI. To capture stable, state-level modulation rather than transient event-related responses, the posterior uncertainty was averaged across all time steps within each sequence and across the 10 selected variables. We then calculated the participant-level mean uncertainty for each attentional condition and compared the difference using a paired t-test.

Subsequently, to derive a single representative index of precision modulation and avoid the multiple comparisons problem in subsequent correlation analyses, we applied Principal Component Analysis (PCA) to the condition differences (*σ*_*SF*_ − *σ*_*BF*_) of the 10 individual BBI variables. The first principal component (PC1) explained most of the variance (67.2%, compared to approximately 14.6% for PC2), indicating a single dominant pattern of precision modulation across the network. We therefore extracted the PC1 score for each participant as a representative index of individual precision modulation.

To evaluate the external validity of this model-derived representation, we examined the associations between the individual PC1 scores and psychological, clinical, and behavioral traits. The indices included objective interoceptive accuracy measured by the heartbeat counting task (Garfinkel et al., 2015; Legrand et al., 2022), as well as specific subjective measures. For psychiatric traits, we assessed depressive symptoms using the Beck Depression Inventory (BDI) (Beck et al., 1996; Kojima et al., 2002), trait anxiety using the trait subscale of the State-Trait Anxiety Inventory (STAI-Trait) (Hidano et al., 2000; Spielberger et al., 1971) to capture stable individual characteristics, and rumination using strictly the rumination subscale of the Rumination-Reflection Questionnaire (RRQ) (Takano & Tanno, 2008; Trapnell & Campbell, 1999). For subjective interoceptive awareness, we used the Multidimensional Assessment of Interoceptive Awareness (MAIA) (Fujino, 2019; Mehling et al., 2012). Based on an a priori hypothesis focusing on attentional control during the BF condition, we selected five specific MAIA subscales (Noticing, Not-Distracting, Attention Regulation, Self-Regulation, and Body Listening) that theoretically reflect top-down and bottom-up attention to bodily sensations.

Given the relatively small sample size (N = 33), we calculated Pearson’s correlation coefficients and assessed their statistical significance using non-parametric permutation testing with 10,000 iterations. To control multiple comparisons, False Discovery Rate (FDR) correction was applied to the permutation-based p-values independently within each targeted trait family. All statistical tests were two-tailed, and the significance level was set at *α* = 0.05.

## Acknowledgements

We are grateful to all the participants in this study.

## Author contributions

[Contribution statement]

## Data and code availability

The data is not publicly available because we did not obtain consent from the participants.

