## Supplementary Fig. 1-2, Supplementary, Supplementary Table1, Fig. 2, Supplementary Fig. 1, Supplementary Fig. 3 for "A hierarchical generative model reveals enhanced latent precision of brain-body interaction dynamics during interoceptive attention"

Supplementary file for Shinagawa *et al*., “A Generative Model of Multimodal Physiological Data Captures Brain–Body Interactions Modulated by Interoceptive Focus”

Supplementary Figures:

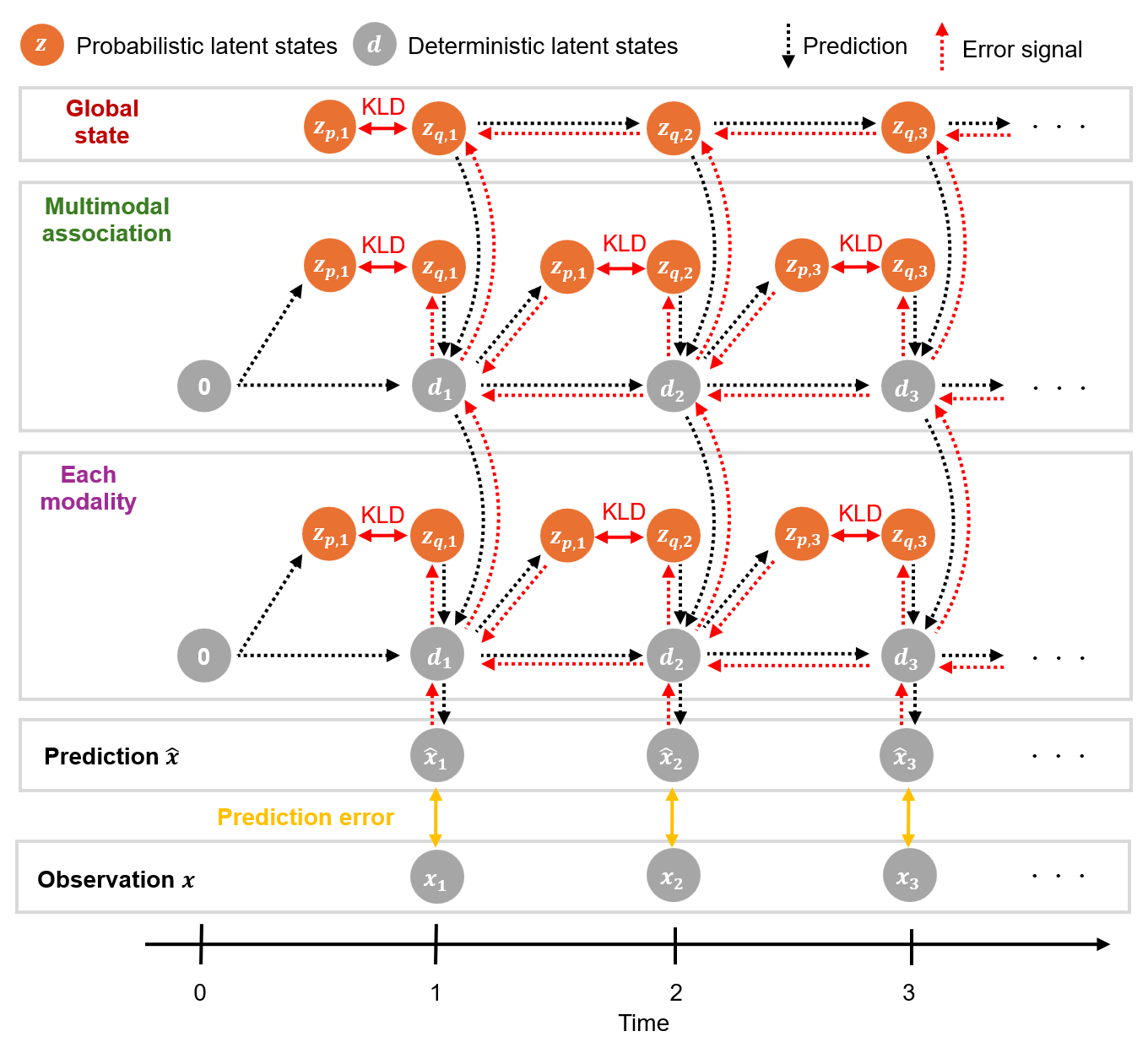

**Supplementary Fig. 1:** **Temporal processing of the V-RNN.**

The network optimizes the posteriors of latent variables 𝑧_1:𝑇 across all modules and updates time-constant synaptic weights by minimizing the free energy F over the duration 𝑇 of training sequences of each modality. Initial deterministic states of all modules are set to zero. For simplicity, only one modules is depicted for each modality module. Abbreviation: KLD, Kullback-Leibler divergence between posterior and prior latent variables.

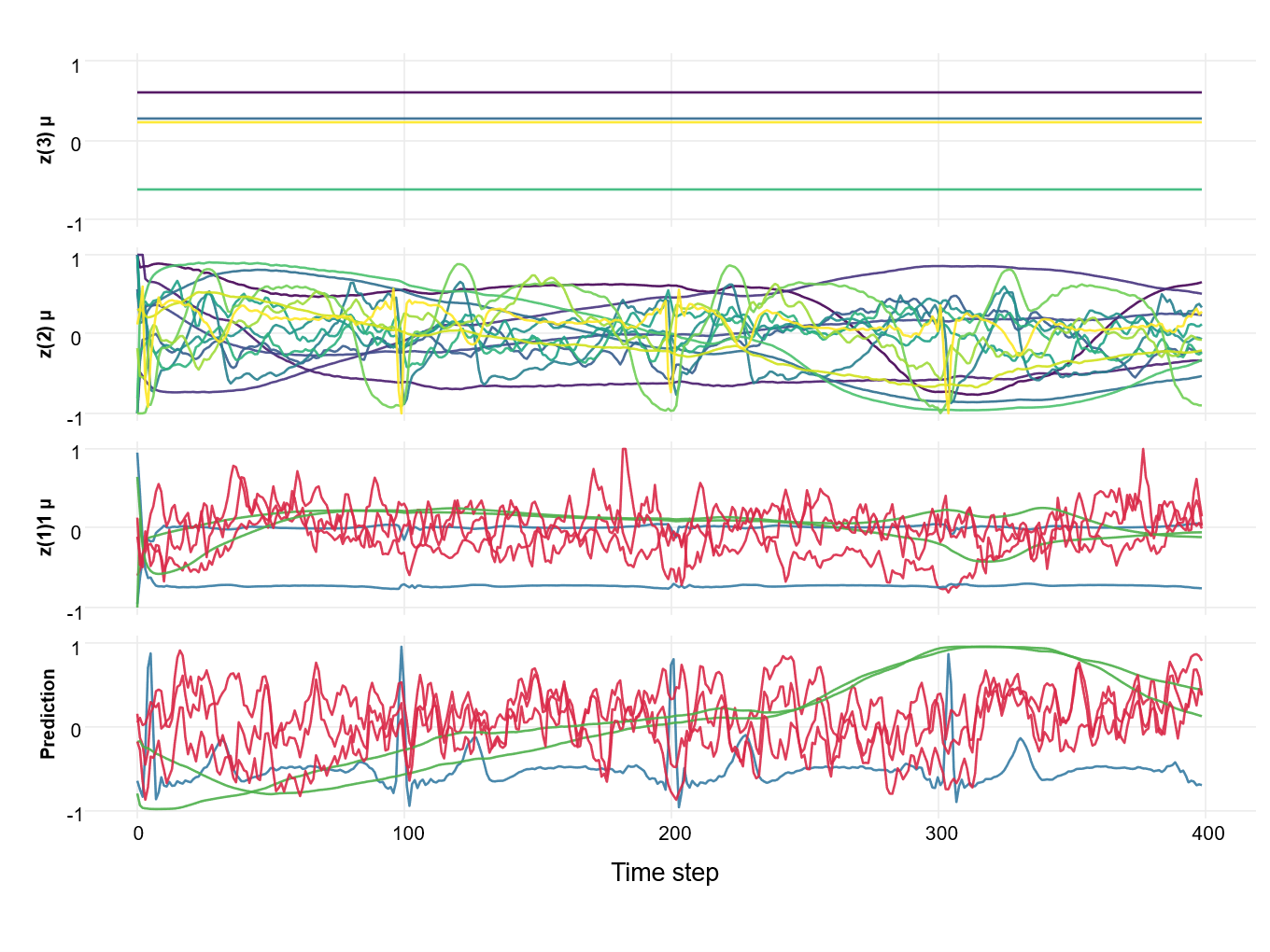

**Supplementary Fig. 2:** **Latent state dynamics of each layer.**

Group Prediction and latent state sequences in training phase. This figure illustrates the sequence of predictions and the mean (μ) of posterior distributions of latent states z(1), z(2), z(3), generated in response to observed data. Predicted and observed EEG signals are visualized by selecting 3 channels from the original channels for visual clarity. Similarly, the z(1)µ sequences were also visualized by selecting 3 latent.

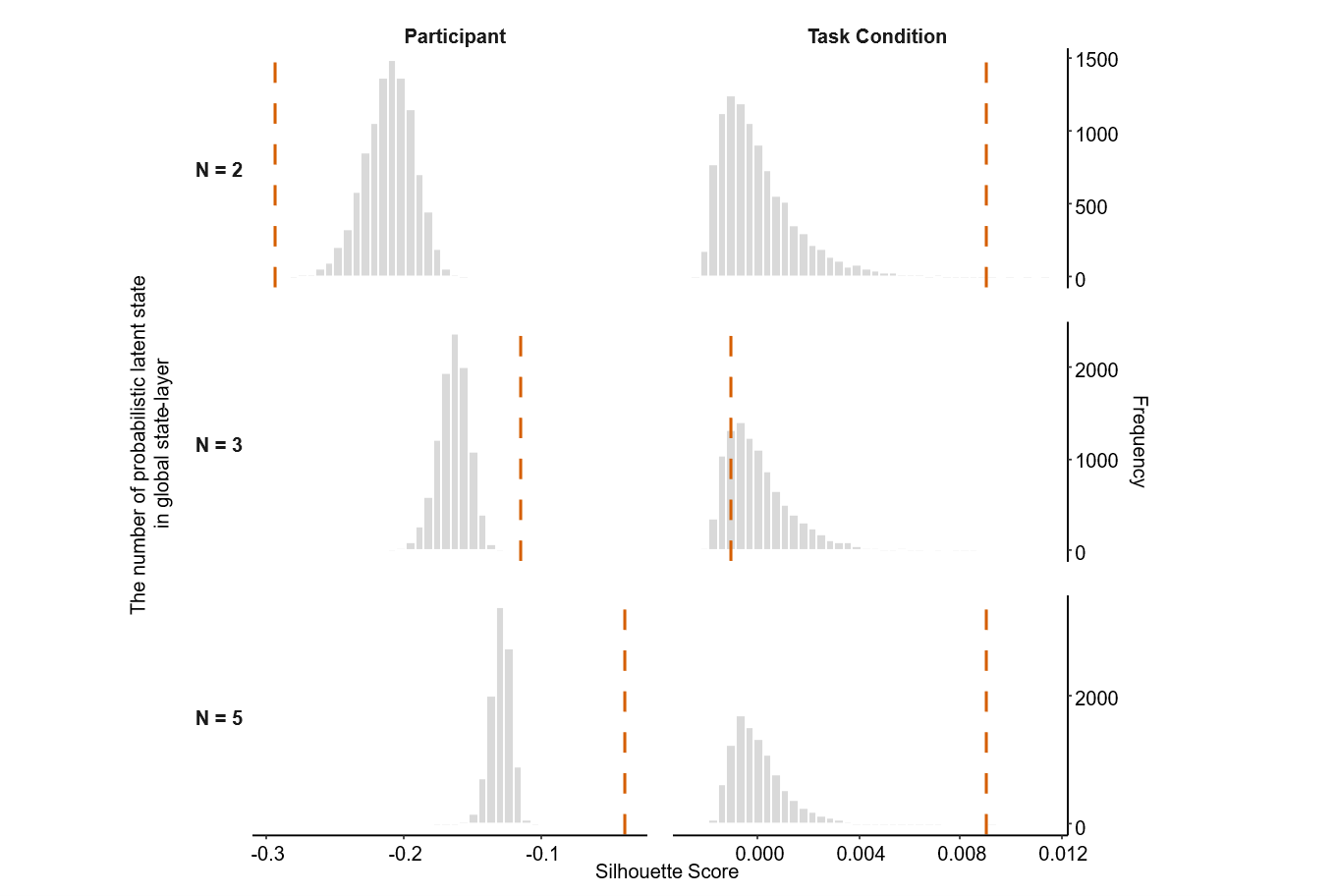

**Supplementary Fig. 3:** **Sensitivity of global state-layer dimensionality.**

To assess robustness of higher-level representations, we repeated the latent-space analysis while varying the number of probabilistic latent state in global state-layer (N=2–5). For each N, to quantify organization in the original executive latent space, we computed silhouette scores for condition and identity and evaluated significance against permutation-based null distributions obtained by randomly shuffling the corresponding labels. Across global state dimensionalities, the structure of the latent space depended on N: for N< 4, organization was significant for either attentional condition or participant identity, whereas for N ≥ 4 the global state latents exhibited above-chance organization for both factors, consistent with a jointly self-organized representation of state-like and trait-like information. Observed silhouette scores (vertical lines) are shown relative to permutation-based null distributions (histograms), and p-values are reported for each N.

Supplementary Table:

**Supplementary Table1: Correlations between the association-layer uncertainty modulation index and questionnaire measures**

| **Variables** | **r** | **p** |
| --- | --- | --- |
| Anxiety | -0.324 | 0.097 |
| BDI | -0.021 | 0.906 |
| HBC | -0.142 | 0.448 |
| MAIA_1 | 0.126 | 0.619 |
| MAIA_2 | -0.414 | 0.046 |
| MAIA_4 | 0.316 | 0.128 |
| MAIA_6 | 0.432 | 0.046 |
| MAIA_7 | 0.051 | 0.778 |
| RRQ | -0.451 | 0.033 |
